# Refinement of Nucleus Accumbens Neuronal Dynamics During Cocaine Self-Administration Training

**DOI:** 10.64898/2026.02.26.708295

**Authors:** Linjie Jin, Xiguang Qi, Jianwei Liu, William J. Wright, Terra A. Schall, King-Lun Li, Bo Zeng, Charles Wang, Lirong Wang, Yan Dong

## Abstract

Drug addiction is an acquired motivational-behavioral state that begins with drug taking, which is comprised of a series of phases, including initial acquisition, stabilization, habituation, and maintenance. In rodent models of cocaine self-administration, the forebrain region nucleus accumbens (NAc) has been critically implicated in the acquisition-maintenance process of drug taking and seeking behaviors. However, it remains unknown how NAc neurons shift their activity patterns in response to these phasic transitions during cocaine taking. To examine this, we used GCaMP6m-based in vivo Ca^2+^ imaging to monitor activities of principal medium spiny neurons (MSNs) in the NAc across eleven days of cocaine self-administration. Behaviorally, mice exhibited progressive stabilization of operant responding and locomotion across 11 days of cocaine self-administration. During the early training days, we detected a portion of NAc neurons—a potential neuronal ensemble—that exhibited increased activities temporally contingent to the lever-press for cocaine. The number of NAc neurons exhibiting contingent activity increased progressively over the first three training days and then decreased gradually during the later training days, exhibiting expansion-refinement dynamics that may correspond to the acquisition and subsequent stabilization/maintenance of cocaine self-administration. Using a neuron-tracking technique, we found that the lever-press-contingent NAc ensemble exhibited substantial compositional dynamics, with neurons dropping into and out across training days. These activity features of lever-press-contingent neurons may represent key circuit dynamics of the NAc that transition the acquisition toward the maintenance of cocaine-taking behavior.

## Introduction

Substance use disorder (SUD) is an acquired motivational-behavioral state that begins with drug taking (Volkow et al., 2019). In rodent models of cocaine self-administration, the process begins when animals first learn the operant response to obtain the drug—a phase called acquisition—followed by maintenance where the operant response stabilizes and persists (Dong et al., 2017). Although these behavioral phases demand distinct neuroeconomic and cognitive efforts, the critical supporting evidence detailing the shift in neural dynamics underlying these phases transitions is lacking (Bickel et al., 2011; Feldstein Ewing and Chung, 2013). To gain insight, we focused on the nucleus accumbens (NAc), a key forebrain region critically implicated in regulating both the acquisition and maintenance of drug taking behavior.

Medium spiny neurons (MSNs) are the principal neurons in the NAc, comprising >90% of its neuronal population (Meredith et al., 2008). These MSNs receive convergent glutamatergic inputs from various limbic and paralimbic brain regions that process reward information (Kelley, 2004). Lacking intrinsic pace-making mechanisms, action potential firing in MSNs is primarily driven by these glutamatergic synaptic inputs, gated by channel-based membrane responsiveness (Wright and Dong, 2020, 2021). In awake rodents, NAc MSNs reside primarily in a functionally active state (the upstate), where they fire action potentials continuously (Mahon et al., 2006). The acute delivery of drug or nondrug rewards (unconditioned stimuli) instantly increases action potential firings of a specific subset of NAc MSNs, forming a reward ensemble (Carelli, 2002). In GCaMP-based Ca^2+^ imaging studies of individual neurons, increased intensities of GCaMP fluorescence predict increased action potential firings (Ali and Kwan, 2020). In mice well-trained with operant responses, an ensemble of NAc MSNs exhibits heightened Ca^2+^ activity around the act of reward taking (Chalhoub et al., 2024; Clarke et al., 2024; Schall et al., 2024; Domingues et al., 2025). These temporal-dynamics of NAc neurons may thus represent key activity patterns that continuously evolve to support the different phases of reward taking behaviors.

Repeated exposure to cocaine or other drugs of abuse induces adaptive changes in both glutamatergic synaptic inputs and membrane properties, which potentially alter the firing patterns of NAc MSNs across periods of drug taking (Wright and Dong, 2020). In this study, we sought to understand how the NAc neurons associated with cocaine taking evolve over the course of self-administration training. Using GCaMP-based in vivo Ca^2+^ imaging, we detected a population of NAc neurons with increased activities temporally contingent to the act of cocaine taking on each training day, thus forming a lever-press neuronal ensemble. The number of neurons in this ensemble increased over the initial training days and then declined over the following days toward the level observed on training day 1. These expansion-refinement dynamics may correspond to the acquisition and subsequent stabilization/maintenance of cocaine self-administration. Furthermore, our tracking analysis revealed that this NAc ensemble exhibited substantial compositional dynamics, with individual neurons dropping into and out of the active cohort across training days. These features of the lever-press ensemble may represent key circuit dynamics of the NAc that contribute to the behavioral transition from the acquisition toward the maintenance of cocaine-taking behavior.

## Methods and Experimental Procedures

### Subjects

Seven male wildtype C57BL/C mice from the Jackson Laboratories were used for cocaine self-administration training. Eight D1-Cre and eleven D2-Cre C57BL/C male mice were used for sucrose self-administration training. We used male mice in this study. This choice was based on preliminary results, which showed that, under our experimental conditions, male mice exhibited substantially higher tolerance to the head-equipped miniscope/GRIN lens assembly. With the head stage, male mice provided a more stable operant response during self-administration compared to female mice, minimizing potential variability in data collection and analysis. At the beginning of the experiments, mice were ∼7 weeks old (∼24 g), housed on a 12-h light/dark cycle (light on/off at 7:00/19:00) with food and water available ad libitum. All mice were used in accordance with protocols approved by the Institutional Care and Use Committees at the University of Pittsburgh.

### In vivo virus injection and GRIN Lens Implantation

To monitor the activity of NAc neurons in vivo, we unilaterally infused an AAV9 vector expressing GCaMP6m in the NAc of the right hemisphere via stereotaxic microinjection. AAV9 we used was pAAV.hsyn.GCaMP6m.WPRE.SV40 (titer ≥ 2.7 x 10^13^ GC/ml), purchased from Addgene (Watertown, MA).

Mice were anesthetized with a xylazine–ketamine mixture (5–10/50–100 mg/kg, i.p.) and received carprofen (5 mg/kg, s.c.) for additional analgesia. After exposing the skull, a ∼1-mm diameter hole was drilled above the NAc. A 32-gauge microinjection needle was lowered to the following stereotaxic coordinates (in mm: AP +1.50; ML +0.73; DV –4.25). A 0.5-µL viral solution was delivered at a rate of 200 nL/min, and the needle was left in place for an additional 5 min to minimize backflow. During the same surgical procedure, a 0.5-mm Gradient-Index (GRIN) lens (Grintech) was implanted directly above the NAc. After a minimum of 4 weeks for viral expression, the miniscope baseplate was attached, and expression was assessed prior to behavioral training. The above coordinates that we used for virus injection and GRIN lens placement targeted the NAc shell. Our post hoc histological examination confirmed that the lens positions were centered toward the NAc shell.

### Catheter implantation

Cather implantation for self-administration in mice was performed as described previously (Yu et al., 2017; Yu et al., 2022; He et al., 2023a; He et al., 2023b). Briefly, mice were anesthetized with a xylazine-ketamine mixture (5-10/50-100 mg/kg, i.p.). A silicone catheter, constructed from silastic tubing (length ∼5 cm, inner diameter 0.012 in, outer diameter 0.025 in) was inserted into the jugular vein, passed subcutaneously to the midscapular region, and connected to a vascular access button (Instech Laboratories) implanted under the skin. Mice were allowed 5-7 days for recovery following surgery. Catheters were flushed every 24 h during the recovery and training periods with sterile saline containing cefazolin (3 mg/mL) and heparin (3 units/mL).

### Cocaine self-administration

Cocaine self-administration was conducted in operant-conditioning chambers (29.53 × 24.84 × 18.67 cm^3^) enclosed within sound-attenuating cabinets (Med Associates). Each chamber was equipped with an active and an inactive lever, a food dispenser, a conditioned stimulus (CS) light at each lever-press hole, a house light, and a speaker for auditory cues. No food or water was provided in the chamber during training or testing sessions.

On day 1, mice underwent a single overnight session (∼12 h) under a fixed-ratio 1 (FR1) schedule. An active lever-press resulted in a cocaine infusion (0.75 mg/kg over 3–6 s), paired with illumination of the CS light, the house light, and an auditory cue (1,000 Hz, 80 dB, 20 pulses at 1 Hz). The light and auditory cues remained on for 20 s, during which additional lever-presses were recorded but did not produce additional cocaine infusions. After the 20-s timeout, all cues were terminated, and the next active lever-press again resulted in cocaine infusion. Presses on the inactive lever were recorded but had no programmed consequences. Only mice that obtained a minimum of 80 cocaine infusions during the overnight session were advanced to the subsequent 11-day cocaine self-administration procedure, as previously described (Yu et al., 2017; Yu et al., 2022; He et al., 2023a; He et al., 2023b).

During the 11-day training period, mice completed daily 2-h sessions using the same FR1 procedure. These daily sessions were conducted within the 10:00 – 14:00 time window across the entire 11-day training procedure to reduce circadian-related variability. GCaMP6m-based Ca^2+^ imaging was performed every other day (d1, d3, d5, d7, d9, d11). Mice were briefly head-restrained for miniscope installation prior to self-administration session (UCLA v3; wire-free version, Open Ephys Production). Imaging recordings were restricted to the first 20 min of each session to minimize photobleaching and accommodate battery capacity. On non-imaging days, mice were trained while fitted with a dummy scope matched in weight and dimensions. Mice that failed to meet the final self-administration criteria (≥15 infusions per session, 70% active-to-inactive lever-press ratio) were excluded from further analysis.

### Sucrose self-administration

Sucrose self-administration was conducted in the same operant chamber used for cocaine self-administration, except additionally equipped with a sucrose sipper. The training procedure consisted of a 12-h overnight session followed by a 10-day training phase, during which the lever-press led to an extraction of sucrose sipper delivering 0.1 mL of 10% sucrose solution and a 10-s presentation of sound-light compound cues. This was followed by a 10-s timeout period, during which additional active lever-press was recorded but not reinforced. Detailed procedure was described in our previous studies (Schall et al., 2024).

### DLC-based calculation of movement

Animal movement was analyzed using the DeepLabCut (DLC) algorithm (Mathis et al., 2018; Nath et al., 2019). Ten individual body parts were manually labeled for 20 frames, after which their coordinates were automatically tracked by DLC. For each self-administration session, a 20-minute video was recorded. The main DLC parameters were configured as: the augmenter-type was set to imgaug, and a ResNet-152 network pre-trained on ImageNet was used. All other parameters were kept at the default DLC values. For movement quantification, the “skull top” label was used to represent the mouse’s position. The DLC extracted positions were down sampled to 5 Hz by selecting the coordinates with the highest likelihood within every 0.5-s or 0.2-s window from the original 15–30 Hz recordings for distance or rotation analyses. Coordinates with likelihood values below 0.7 were discarded and then interpolated. Pixel coordinates from all videos were normalized to a 1920 × 1080 frame, and the chamber area was divided into grids of 50 × 50 normalized pixels for subsequent analysis.

To match the time course of in vivo Ca^2+^ imaging, our analysis focused on the first 20 min of each training session. We calculated the total travel distance, the accumulated weighted rotation, and the area-to-distance ratio to quantify mice’s locomotion within the chamber. The sample for distance calculation was further down-sampled to 2 Hz to minimize the path length inflation caused by DLC estimation errors. Similar results were obtained using 5-Hz down-sampling. The accumulated weighted rotation was calculated across all −5 s to +5 s time windows relative to each lever-press, with non-reinforced lever-press excluded. The sample of accumulated weighted rotation remained at 5 Hz (i.e., 0.2-s intervals), because the clockwise versus counterclockwise rotations carry opposite signs, allowing estimation errors to cancel out. For each interval [t_i_, t_i+1_J, the mouse’s coordinates at the beginning (x_i_, y_i_), and the end (x_i+1_, y_i+1_) were determined. The center of the mouse’s entire trajectory during the session was defined as the midpoint (x_C_, y_C_). The original rotation angle e_i_ was computed as the angle between the two position vectors from the midpoint to the start and end coordinates: 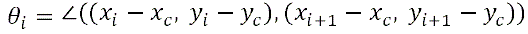. The weighted rotational angles for each interval was defined as: 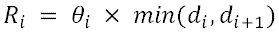, where 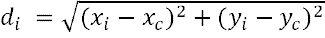, and 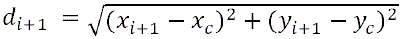. Clockwise and counterclockwise rotations were assigned with the opposite signs, and accumulated weighted rotation was calculated the sum of all R_i_. To quantify the variability of trajectories when the mouse approached and exited the active lever across trials, we defined the area-to-distance ratio. The distance was calculated across all −5 s to +5 s time windows relative to each lever-press, with non-reinforced lever-press excluded. The area was estimated by the number of 50 × 50-pixel grids that the mouse passed through at least once within the −5 to +5 s window around each lever-press.

### Analysis of Ca^2+^ signals

*Raw video data* (∼320 × 320 μm, 20 Hz) from each imaging session were processed *using the Python toolbox CaimAn* (Giovannucci et al., 2019). We applied an established pipeline (Zhou et al., 2018) for 1-photon imaging, which consists of the non-rigid motion correction (NoRMCorre) (Pnevmatikakis and Giovannucci, 2017) algorithm as well as constrained non-negative matrix factorization endoscopic (CNMF-E) source separation and quality control metrics to evaluate final datasets (Pnevmatikakis et al., 2016). For cross-trial normalization, Ca^2+^ traces were divided into 100-ms bins and then subjected to z-score standardization (mean = 0; SD = 1).

### Heatmap and sorting

For all heatmaps, each row represented an individual neuron with trial-averaged z-scores aligned with the lever-press timepoint. To obtain the trial-averaged z-scores in each neuron, we extracted the z-scores of all activities of this neuron across the entire 20-min imaging period, and then selected the z-score data across the time window (e.g., 20 s) of all the lever-press trials for averaging. For controls, neurons were aligned with randomly selected, trial number-matched time windows. For sessions with fewer than 10 trials, we used a fixed set of 10 random windows to minimize variability due to limited sampling. Visualization details varied among figures to show specific aspects of the data. For example, In **Figure 2f**, neurons were sorted by the order in which they reached their peak Ca^2+^ activities, while in **Figure 3a-f**, neurons were sorted by their mean activities within the 0-5-s post-lever-press time window. All heatmaps were generated using Python package *seaborn* (Waskom, 2021) and *matplotlib* (Hunter, 2007).

**Figure 1.**
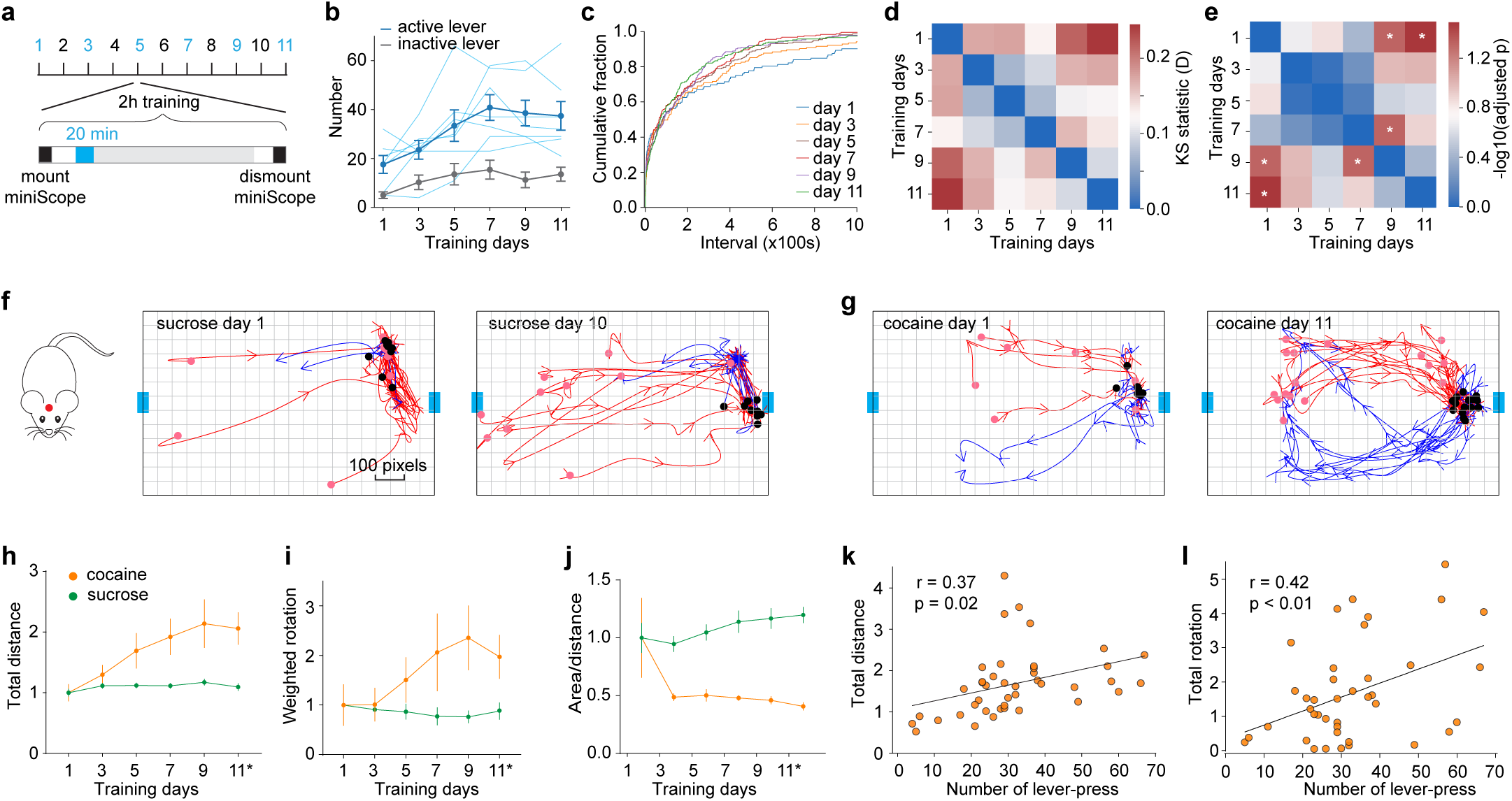
Progressive stabilization of operant responding and locomotion during cocaine SA training. (**a**) Schematic showing the 11-day cocaine self-administration procedure and the periods during training sessions when in vivo imaging was performed. (**b**) Summaries of operant responses during cocaine self-administration training in the seven mice used for in vivo imaging. Light blue lines indicate data from individual mice [F_5_, 30 = 6.6, p < 0.01, main effect of day; F_1,6_ = 26.4, p < 0.01, main effect of condition (active lever-press vs. inactive lever-press), two-way ANOVA repeated measures]. Note: One mouse was excluded from the analyses in Fig. 2g, h and Fig. 3 due to the missing video/imaging data on a training day. (**c**) Cumulative distribution of the inter-press interval, defined as the time between two consecutive lever-presses. (**d**,**e**) Heatmaps showing results from the Pairwise Kolmogorov–Smirnov (KS) test: KS statistics (D) of cumulative distribution of inter-press intervals between two self-administration days (**d**), and the (–log_10_)-adjusted p-values after false discovery rate (FDR) correction (**e**); higher values suggesting larger differences (day 1 vs. day 9, p < 0.05; day 1 vs. day 11, p = 0.03, day 7 vs. day 9, p < 0.05). (**f**) Diagram (left) showing the position of the “skull top” label (red dot) used for tracking and movement trajectories of an example sucrose-trained mouse during the approach to and exit from active lever on days 1 and 10. Red and blue arrows indicate trajectories 5 s before versus after each lever-press, respectively. Pink and black dots mark the mouse’s position 5 s before and the timepoint of the lever-press, respectively. Blue squares mark the positions of active (right) and inactive (left) levers. Grids were used to calculate the area-to-distance ratio shown in (**j**). (**g**) Movement trajectories of an example cocaine-trained mouse on days 1 and 11. (**h**) Summaries showing that the total movement distance gradually increased over days of cocaine self-administration (F_1,24_ = 15.24, p < 0.01, main effect of group; F_5,111_ = 16.89, p < 0.01, main effect of day; F_5,111_ = 11.37, p < 0.01, day x group; two-way ANOVA mixed effect model). (**i**) Summaries showing that the accumulated weighted rotations gradually increased over days of cocaine self-administration (F_1,24_ = 8.30, p < 0.01, main effect of group; F_5,110_ = 2.21, n.s., main effect of day; F_5,110_ = 3.93, p < 0.01, day x group; two-way ANOVA mixed effect model). (**j**) Summaries showing that the area-to-distance ratio during the –5 s to 5 s lever-press time window decreased over days of cocaine self-administration. Day 11* indicates training day 10 for sucrose mice. (F_1,24_ = 45.58, p < 0.01, main effect of group; F_5,110_ = 1.27, n.s., main effect of day; F_5,110_ = 2.95, p = 0.02, day x group; two-way ANOVA mixed effect model) (**k**) Summaries showing a correlation between the total movement distances and lever-press numbers in cocaine mice (Pearson’s r = 0.37, p = 0.02). (**l**) Summaries showing a correlation between the total accumulated rotations and lever-press numbers in cocaine mice (Pearson’s r = 0.42, p < 0.01).

**Figure 2.**
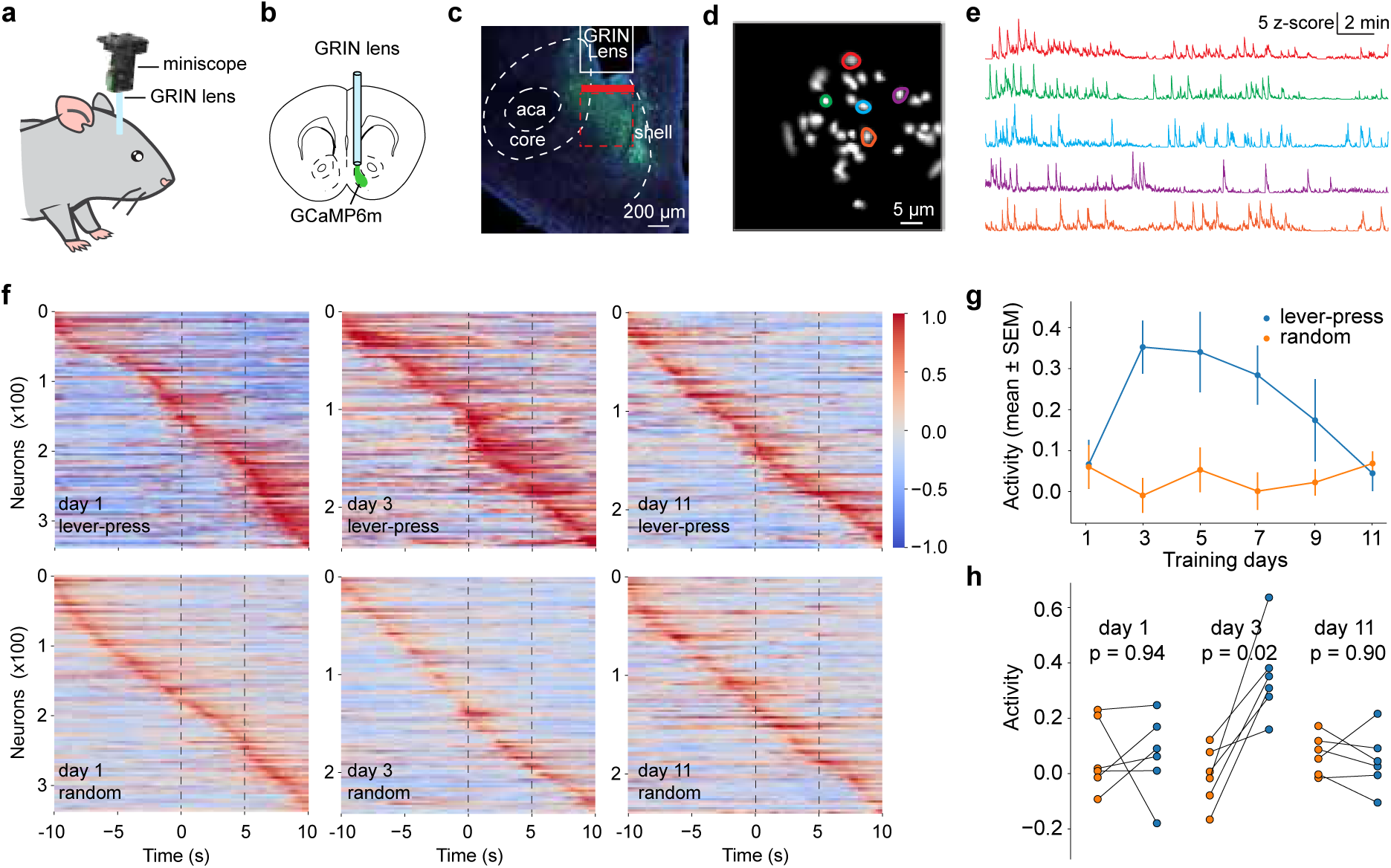
Ensemble activities of NAc neurons exhibited bell-shaped changes over 11 days of cocaine self-administration. (**a**,**b**) Diagrams showing the installation of miniscope and GRIN lens (**a**) and in vivo imaging of Ca^2+^ activities of neurons in the NAc (**b**). (**c**) Image showing GCaMP6 expression in the NAc shell, position of the GRIN lens, the imaging plane relative to the GRIN lens (indicated by horizontal red bar), and the range of imaging planes summarized from the postmortem verification of the mice used for this study (indicated by the red dashed-line box). (**d**) Example image showing individual neurons with Ca^2+^ activities imaged and extracted using the CNMF-e algorithm. (**e**) Ca^2+^ transients extracted from example neurons in (**d**) (traces and neurons were color matched). (**f**) Heatmaps of neuronal activity during randomly selective versus lever-press time windows on training day1 (left column), day3 (middle column), and day11 (right column). Neurons were sorted by the time of their peak activity within the –10 s to +10 s time window around lever-press. (**g**) Summaries of NAc neuron activity during the 0-5 s time window post-press. Neuronal activity was assessed as trial-averaged z-scores for individual mice (F_5,25_ = 3.9, p < 0.01, day x lever-press/random, two-way ANOVA repeated measures) (i) Comparison of trial-averaged NAc neuron activity in individual mice between the randomized and lever-press 5-s time window on training days 1, 3 and 11 (day 1, t_5_ < 0.1, p = 0.94; day 3, t_5_ = 4.4, p = 0.02; day 11, |t_5_| < 0.1, p = 0.89; paired t-test).

**Figure 3.**
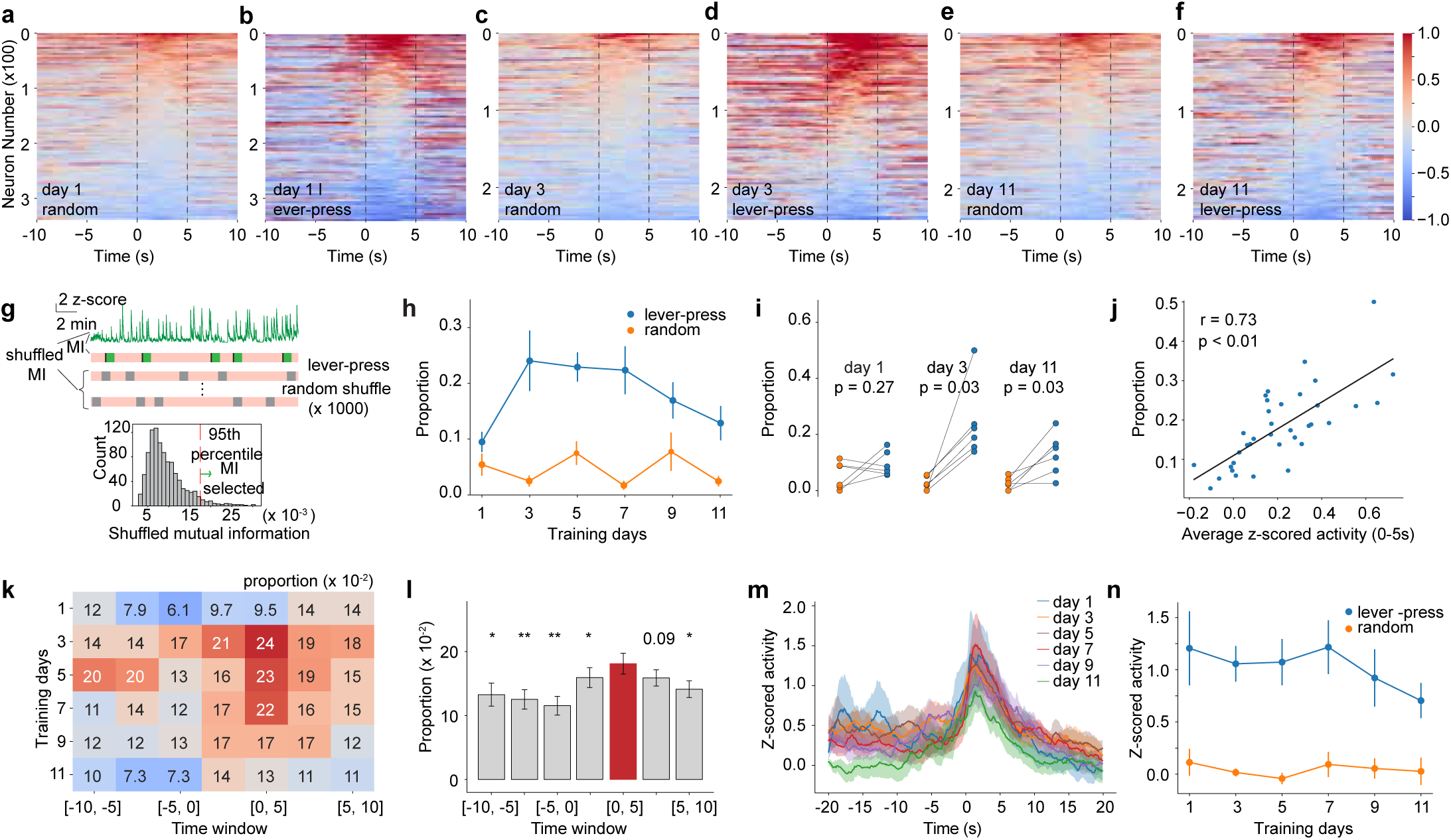
Mutual information analysis reveals bell-shaped changes in the number of lever-press-responsive neurons over 11 days of cocaine self-administration. (**a-f**) Heatmaps showing NAc neuron activity during random versus lever-press 20-s time windows on cocaine self-administration day 1 (**a, b**) day 3 (**c, d**), and day 11 (**e, f**). Neurons were sorted by the trial-averaged z-scores during the 0-5-s time window. (**g**) *upper panel*: Schematics illustrating the strategy of using MI to determine lever-press-responsive neurons based on the temporal contingency of neuronal activity to the behavioral response. *lower panel*: Shuffled MI distribution for an example neuron, generated by randomly shuffling the timing of lever-press 1000 times. Neurons were defined as lever-press-responsive if their MI with lever-press timing exceeded the 95th percentile in the shuffled distribution. (**h**) Summaries showing bell-shaped changes in the proportion of lever-press-responsive neurons among all recorded NAc neurons over the 11-day cocaine self-administration (F_5,25_ = 3.0, p = 0.03, main effect of day; F_1,5_ = 185.3, p < 0.01, lever-press vs. random; F_5,25_ = 2.5, p = 0.05, day x lever-press/random; n = 6, two-way ANOVA repeated measures). (**i**) Comparison of the proportion of lever-press-responsive neurons between randomly selected and lever-press time window on cocaine self-administration days 1, 3, and 11 (day 1, t_5_ =1.2, p = 0.27; day 3, t_5_ = 3.8, p = 0.03; day 11, t_5_ = 3.4, p = 0.03; paired t-test with false discovery rate correction). (**j**) Summaries showing a positive correlation between the averaged activity of all recorded neurons over 0-5 s post-press and the proportion of lever-press-responsive neurons. Each data point representing one mouse on one self-administration day (Pearson’s r = 0.73, p < 0.01). (**k**) Heatmap showing proportions of lever-press-responsive neurons across different time windows versus self-administration days (F_6, 30_ = 3.5, p = 0.01, time window, n = 6, two-way ANOVA repeated measures). (**l**) Summaries showing the proportions of lever-press-responsive neurons across different time windows, with statistical annotations indicating the comparison of the 0–5 s window with other time windows (0–5s vs. others: t_35_ = 2.6, p = 0.02, –10 to –5 s; t_35_ = 4.2, p < 0.01, –7.5 to –2.5 s; t_35_ = 5.4, p < 0.01, –5 to 0 s; t_35_ = 2.2, p = 0.04, –2.5 to 2.5 s; t_35_ = 1.7, p = 0.19, 2.5-7.5 s; t_35_ = 2.3, p = 0.04, 5 to 10 s; t-test). (**m**) Summaries showing averaged activities of lever-press-responsive neurons on self-administration days (n = 6 mice). (**n**) Summaries showing averaged activities of lever-press-responsive neurons (over the 20-s lever-press time window) across training days (F_5,25_ = 1.0, p = 0.45, main effect of day; F_5,25_ = 37.2, p < 0.01, main effect of lever-press vs. random; F_5,25_ = 0.5, p = 0.78, day × lever-press/random; n = 6, two-way ANOVA repeated measures).

### Mutual information and selectivity

The mutual information analysis (MI) (Shannon, 1948) was used to quantitatively select neurons exhibiting activity contingent to lever-pressing. The MI provided a value that reflected the degree of dependency, covering linear and nonlinear relationships, between the two variables: neuronal Ca^2+^ transients and the behavioral array (lever-presses). For computation, the behavioral array was structured with timepoints around the lever-press (0-5 s) set to 1 and all other timepoints set to 0. Ca^2+^ activities were binned using the Freedman–Diaconis estimator, as required for the MI setting. To classify responsiveness, a null distribution was established by randomly shuffling the behavioral array 1000 times for each neuron (**Figure 3g**). Neurons whose observed MI value exceeded the 95^th^ percentile of the null distribution were classified as behaviorally responsive. For statistical analysis, we calculated the ratio of these MI-selected neurons to the total number of extracted neurons per mouse. To assess the temporal selectivity, we evaluated changes in the neuron’s MI by sliding the behavioral time window around the lever-press timepoint (**Figure 3k**). The MI value was calculated using the *mutual_information_score* function from the *Scikit-learn.metrics* package (2011). One mouse was excluded from this analysis due to the absence of lever-presses on training day 1.

### Tracking neurons across sessions

We used a combined process of image alignment, probabilistic modeling, clustering, and final registration (Pnevmatikakis et al., 2016) to track neurons across multiple imaging days. Candidate neuron pairs were selected based on the similarities of their shapes and relative spatial positions (Sheintuch et al., 2017) (**Figure 4a** and **4b**). To account for local deformation between imaging sessions, we employed non-rigid registration with the transformation smoothness set to 1. Candidate pairs exceeding the between-session distance threshold of 15 μm were excluded. The robustness of this spatial registration was validated by testing various combinations of transformation smoothness (e.g., 0.8, 1, 1.2, and 1.5) and distance thresholds (e.g., 10, 15, 20, and 30), which consistently produced substantial overlap in the resulting candidate pairs.

**Figure 4.**
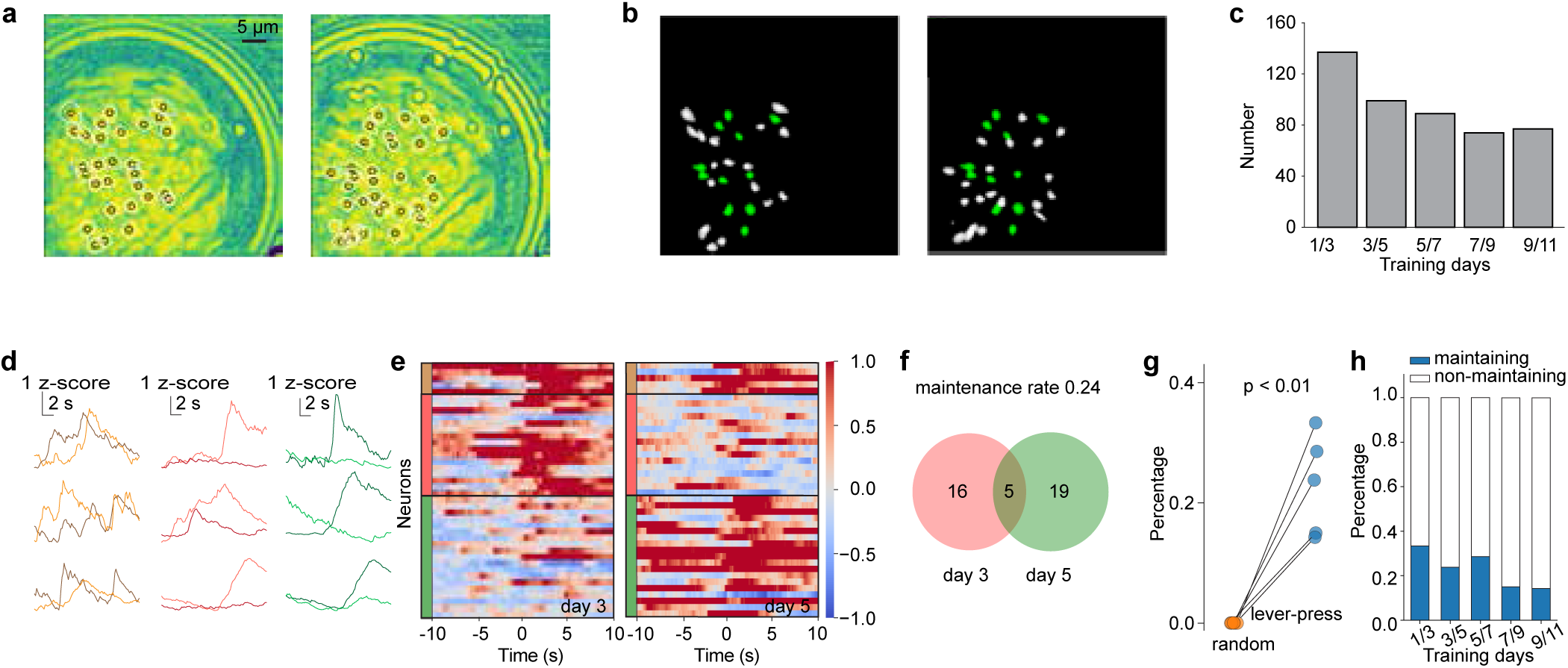
Partial overlap of lever-press-responsive neurons across different days of cocaine self-administration. (**a**) CNMF-e-based mapping and extraction of GCaMP6m-imaged NAc neurons in an example mouse on different days of cocaine self-administration. (**b**) Tracking analysis of images in panel **a** showing matched (green) versus unmatched (white) neurons over the two training days. (**c**) Summaries showing numbers of matched neurons in 7 mice between adjacent cocaine self-administration days. (**d**) Examples showing three types of tracked neurons in their trial-averaged activities (light versus dark colors indicating earlier versus later training days, respectively) over self-administration: responsive on both days (left), nonresponsive on earlier day (middle), and nonresponsive on later day (right). (**e**) Heatmap showing three types of tracked neurons regarding their responses to lever-press on days 3 (left panel) and 5 (right panel): gaining, losing, and maintaining neurons (indicated by pink, green, and brown vertical bars, respectively, n = 7 mice). Each row represents one tracked neuron with its activity during the 20-s lever-press time window on day 3 versus day 5. (**f**) Venn diagram showing partial overlap of lever-press-responsive neurons identified in panel **e**. (**g**) Summaries showing a higher cross-day maintaining rate of NAc neurons responsive to the lever-press timepoint compared to a random timepoint (lever-press, 0.23 ± 0.04; random, 0.00±0.00; t_4_ = 6.16, p < 0.01, paired t-test). Each data point represents the maintaining rate across two imaging days (days 1-3, 3-5, 5-7, 7-9, and 9-11). (**h**) Detailed presentation of the same data in **g**, showing a higher non-maintaining rate of lever-press-responsive neurons compared to cross-day maintaining rate (non-maintaining, 0.77 ± 0.04; maintaining rate, 0.23 ± 0.04; t_4_ = 7.2, p < 0.01, paired t-test).

### Heatmap and division of tracked neurons

To define a neuron as a lever-press-responsive one, we used the same MI-based standard as defined in **Figure 2**. Neurons were categorized into four groups (**Fig. 4d, e**): 1) responsive on both days, 2) responsive only on the later day, 3) responsive only on the earlier day, and 4) unresponsive on both days. The maintenance rate was calculated as the proportion of neurons that were responsive on both days relative to neurons that were only responsive on the earlier day (**Figures 4f-h**).

### Distribution of inter-press intervals

The inter-press interval was defined as the time elapsing between two adjacent lever-presses. Pairwise Kolmogorov–Smirnov (KS) tests were performed, using the *SciPy.stats* (Virtanen et al., 2020) package in Python to assess statistic differences in the distributions across days.

### Drugs and reagents

Cocaine HCl was provided by the NIDA Drug Supply Program, was dissolved in 0.9% NaCl saline for self-administration. Ketamine and xylazine were purchased from a Drug Enforcement Agency-designated vendor at the University of Pittsburgh, and were mixed for use as the anesthetic agent. All other chemicals and reagents were procured from commercial vendors, including Sigma-Aldrich, Tocris, J.T.Baker, Fluka, or Alomone labs.

### Data acquisition and statistics

All results are shown as the mean ± s.e.m. All data collection was randomized. All data sets were consistent with normal distribution based on their Quantile-Quantile plots. All data were analyzed offline, with the statistical significance assessed using two-tailed paired t-test, one-way or two-way ANOVA followed by false discovery rate-corrected post hoc tests. Statistical significance was uniformly set at *p* < 0.05. Statistical analyses were performed using Python package *SciPy.stats* (Virtanen et al., 2020).

## Results

### Progressive stabilization of operant responding over 11-d cocaine self-administration

Cocaine self-administration is a learned behavior that progresses through distinct phases: acquisition, stabilization, and maintenance. To gain insights into such behavioral transitions, we subjected mice to a relatively long-term cocaine self-administration procedure, beginning with a 12-h overnight session followed by 11 consecutive days of 2-h daily sessions (**Fig. 1a**). The results demonstrated a clear and progressive stabilization of the cocaine-taking behavior over this period. Specifically, the number of lever-presses increased progressively across the training days (**Fig. 1b**). Consistent with this change, the inter-press intervals, defined as the time between two effective lever-presses, progressively shifted their distribution toward shorter durations (**Fig. 1c**). Furthermore, the Kolmogorov–Smirnov (KS) test confirmed this increased consistency, revealing a greater similarity in the distribution of inter-press intervals during later training days compared to earlier days (**Fig. 1d, e**). Collectively, these findings confirm an increased stability of the operant response over 11 days of cocaine self-administration.

Accompanying the stabilization of the operant response, the cocaine mice also gradually developed movement patterns that appeared to be drug-specific, which were monitored in comparison with mice trained for sucrose self-administration. At the beginning of the procedure (e.g., training day 1), neither sucrose nor cocaine mice showed a patterned approach trajectory to the active lever, exhibiting either minimal movement near the lever or random exploration within the chamber (**Fig. 1f–g**). However, after days of training (e.g., day 11), the cocaine mice—but not sucrose controls—displayed stereotyped movement patterns. Their entries to and exits from the lever area followed more consistent paths, and they showed increased rotational movements within the chamber (**Fig. 1f–g**).

To quantify these behavioral changes, we analyzed the total movement distance, the accumulated weighted rotation, and the area-to-distance ratio around the lever-press. Across training days, cocaine mice exhibited gradual increases in the total movement distance, suggesting sensitized locomotor activity (**Fig. 1h**). In parallel, cocaine mice exhibited gradual increases in the accumulated weighted rotation, suggesting the formation of a circular movement stereotype after the self-administration was stabilized (**Fig. 1i**). Furthermore, the area-to-distance ratio around the lever-press decreased over training days, suggesting reduced trajectory variability and a more refined, focused approach and exit behavior (**Fig. 1j**). In contrast, neither of these behavioral patterns was not formed in sucrose mice. Importantly, both the total movement distance and the cumulative rotation in cocaine mice were positively correlated with the total number of lever-presses for cocaine (**Fig. 1k–l**). Taken together, these behavioral changes may represent phasic transitions as the cocaine self-administration behavior becomes ingrained and automatic across the 11-day training period.

### Bell-shaped changes of NAc neuronal activity over cocaine self-administration

To visualize the activity of NAc neurons, we virally expressed the Ca^2+^ indicator GCaMP6m in the NAc of mice and installed a gradient-index (GRIN) lens directly above the injection plane (**Fig. 2a, b**). The imaging plane of the GRIN lens was targeted to the NAc shell, which was verified postmortem (**Fig. 2c**). On every other day, we recorded GCaMP6m-manifested Ca^2+^ transients during the first 20 min of the session to assess the activity of individual NAc neurons associated with the operant response (**Fig. 2d-e**).

To assess activity changes, we computed z-scores of Ca^2+^ transients using a 100-ms bin. We defined the time of lever-press as the 0-sec timepoint and examined trial-averaged activities within a 20-s time window (−10 s to +10 s). Sorting NAc neurons by the time they exhibited peak activity revealed that the overall activity intensity increased across self-administration days compared to control data sets, which sampled neuronal activities during randomly selected, trial number-matched 20-s time windows (**Fig. 2f**). In the control time windows, neurons were distributed evenly without apparent aggregation, suggesting that the observed activity pattern around the level-press was temporally specific and not due to random firing. Furthermore, a portion of neurons exhibited aggregated activity increases around the lever-press timepoint, which could be visualized on day 3 (**Fig. 2f**). Indeed, this aggregation, or ensemble activity, peaked on days 3 and 5, with increased activity enriched in the 0-5-s post-press time window, before subsequently declining (**Fig. 2g**).

For quantitative analysis, we calculated the mean activity of all recorded neurons during the 0-5-s post-press time window, using neuronal activities during a randomly selected 5-s window as controls. The lever-press-associated neuronal activity was comparable to random periods on day 1, followed by a marked elevation by day 3. Activity remained elevated through day 5, after which it progressively declined on days 7 and 9, ultimately returning to control levels (**Fig. 2g, h**). This temporal profile resembles a bell-shaped or inverted U-shaped curve, indicating the dynamic changes of population activity in the NAc during the behavioral transition from earlier (e.g., acquisition) to later (e.g., stabilization or maintenance) phases of cocaine self-administration.

### Time-dependent bell-shaped changes of lever-press-responsive neurons

The above results established the 0-5-s time window as a critical period for NAc population activity. To investigate this further, we sorted all recorded neurons with their mean trial-averaged z-scores within this 5-s period, using data from randomly selected non-lever-press periods as controls. The sorting revealed a cluster of high-activity neurons on each self-administration day (Data on days 1, 3, and 11 are shown; **Fig. 3a-f**). Visually, the intensity and size of this cluster were more prominent on days 3 and 5 compared to days 1 or 11. This cluster may represent a lever-press-responsive NAc neuronal population that underwent dynamic changes across the early versus late days of cocaine self-administration.

To define the lever-press-responsive neuronal cluster, we used the Mutual Information analysis (MI). MI quantifies the degree of dependency, both linear and nonlinear, between two variables—in our cases, neuronal activities and lever-presses. For each neuron, we calculated its observed MI value and established a null distribution by randomly shuffling the lever-press timings 1000 times. A neuron was classified as lever-press-responsive if its observed MI value exceeded the 95th percentile of its corresponding null distribution. We then assessed the temporal specificity of lever-press-responsive neurons by sliding a 5-s window (with a 2.5-s increment) across the 20-s period (−10 s – +10 s). Across all time windows, the highest proportion of lever-press-responsive neurons was consistently observed in the 0-5-s post-press window on every training day (**Fig. 3k**). Quantifying the proportion of lever-press-responsive neurons within this 0-5-s time window showed a relatively low level on day 1, a high level through days 3 and 7, and then a decline to low levels by days 9 and 11 **(Fig. 3h, i)**. This day-dependent change mirrors the bell-shaped curve observed for the population-wise mean activity (**Fig 2g, h**). Furthermore, the mean activity of all recorded neurons was positively correlated with the portion of lever-press-responsive neurons during the 0-5-s time window (**Fig. 3j**). Thus, the lever-press-responsive neurons may represent the overall neuronal population for the functional output of NAc during the operant response for cocaine.

In contrast to the dynamic change in the number of responsive neurons, the mean activity intensity of these MI-selected neurons remained stable. Specifically, the trial-averaged z-scores of these responsive neurons were similar during a 40-s lever-press period (**Fig. 3m**) as well as within the 0-5-s time window (**Fig. 3n**) across all training days. Thus, the circuit’s dynamic evolution—the bell-shaped profile—is driven primarily by the recruitment and refinement of the lever-press-responsive neuron cluster, rather than by changes in the activity intensity of individual, constituent neurons.

It has long been hypothesized that NAc neurons are organized in distinct ensembles to mediate different aspects of motivated behaviors (Pennartz et al., 1994; Coss et al., 2022; Pedersen et al.; Arroyo et al., 2024; Pedersen et al., 2024). Lever-press-responsive neurons identified above may represent such an ensemble implicated in cocaine taking. The day-dependent changes in the size of this ensemble—i.e., the number of ensemble neurons, may reflect the differential requirement of the circuit resources during acquisition versus stabilization-maintenance of the cocaine-seeking behavior.

### Dynamic constitution of the lever-press-responsive ensemble

Strong empirical evidence suggests that neuronal ensembles or engrams are constituted by a relatively fixed set of neurons that persist throughout post-learning periods (Josselyn et al., 2015; Tonegawa et al., 2018; Whitaker and Hope, 2018). To test whether the lever-press ensemble adhered to this principle, we tracked NAc neurons across training sessions based on their shapes, relative spatial positions, and Ca^2+^ activity patterns (**Fig.4a, b**). Due to substantial technical challenges and the strict criteria of our 2-step tracking strategy (see **Methods**), we restricted our quantitative analysis to tracking between two adjacent days. As a result, we tracked 137 neurons between days 1 and 3, 99 neurons between days 3 and 5, 89 neurons between days 5 and 7, 74 neurons between days 7 and 9, and 77 neurons between days 9 and 11 (**Fig. 4c**).

Based on the lever-press responsiveness within the 0-5-s time window, applying a MI-selection, we categorized tracked neurons into four groups (**Fig. 4d, e**): 1) maintaining responsive on both days, 2) responsive only on the later day, 3) responsive only on the earlier day, and 4) unresponsive on both days. We found that the combined number of neurons entering and dropping out from the ensemble was larger than the number of maintaining neurons (**Fig. 4f, h**). Furthermore, the maintenance rate of responsive neurons—defined as the proportion of responsive neurons relative to neurons responsive on the earlier day—was consistently low across all tracked pairs (0.23 ± 0.04), yet it was significantly higher than randomized controls (**Fig. 4g**). These results demonstrate substantial compositional dynamics, with neurons continuously entering and leaving the ensemble throughout the course of cocaine self-administration training.

## Discussion

In mice performing reward-trained operant responses, select populations of NAc neurons exhibit activity increases temporally correlated to reward seeking and taking (Schall et al., 2024). Our current study detected such a population of NAc neurons during cocaine self-administration, representing a functional ensemble contributing to cocaine taking. Further analysis indicated that the size of this ensemble rapidly expanded during earlier training days but then declined over later days, suggesting an initial recruitment phase followed by a subsequent refinement phase that aligns with the acquisition and stabilization/maintenance of cocaine self-administration. Importantly, this cocaine seeking ensemble is not compositionally stable: ∼50% of neurons rotate in and out across days. These results depict an ever-evolving feature of a potential NAc ensemble, which may encode the progression of behavioral states in developing and maintaining drug-taking behaviors.

### Cocaine-induced behavioral dynamics

Across the 11-day cocaine self-administration, mice exhibited progressive increases in total movement distance, the development of rotational behavioral patterns, and the stabilization of the approach and exit trajectories around the active lever. The heightened locomotor activity is consistent with the phenomenon of cocaine-induced locomotor sensitization, which reflects an elevated basal psychomotor state, but not likely associated with the lever-press-contingent ensemble activity (Robinson, 1993; Tolliver and Carney, 1994; Schlussman, 1998; Wolf, 1998, 1999; Kalivas et al., 2009). On the other hand, the approaching and exiting movements are two essential components of the entire cocaine-taking behavioral cascade, occurring immediately before versus after the act of lever-press. As such, we speculate that their underlying circuit dynamics are closely linked to the press-contingent ensemble—possibly a pre-press and a post-press ensemble, which, together with the press ensemble, activate sequentially to support the cocaine-taking behavioral sequence. Furthermore, the formation of patterned movements may reflect a transition of the cocaine-taking from an exploratory-acquiring to a stereotypic-habitual behavioral state. In this case, the rotational pattern reached to and plateaued at the peak after 7-9 days of cocaine self-administration, suggesting days 7-9 as a potential timepoint around which mice transitioned from the acquisition to later phases (e.g., stabilization or habituation) of cocaine-taking behavior.

### Formation of the lever-press ensemble

NAc MSNs are thought to be organized into different neuronal ensembles, each encoding a specific aspect of behavior (Pennartz et al., 1994; Robinson and Carelli, 2008; Warren et al., 2017; Wright and Dong, 2021). Constituting >90% of the neuronal population in the NAc (Meredith et al., 2008), MSNs are expected to be the dominant neurons that were sampled through Ca^2+^ imaging and thus the primary contributors to the lever-press–responsive ensemble.

There are two dominant subtypes of MSNs in the NAc, one expressing dopamine D1 receptors (D1R), and the other expressing D2R, with a small portion of MSNs expressing both D1R and D2R (Wong et al., 1999; Lobo and Nestler, 2011; Zinsmaier et al., 2022; Gayden et al., 2023; Bonnavion et al., 2024). Using reporter mice, previous studies show that both D1 and D2 MSNs exhibited time-locked activity increases in response to reward delivery (Coss et al., 2022; Chalhoub et al., 2024; Schall et al., 2024; Domingues et al., 2025). While our current study used wildtype mice without distinguishing neuronal types, it is a reasonable assumption that the lever-press-responsive ensemble is comprised of both D1 and D2 MSNs.

As proposed previously (Schall et al., 2024), although included in the same lever-press ensemble, D1 and D2 MSNs regulate distinct circuit targets. Related to this, the lever-press ensemble was defined by the increased neuronal activity during the 0-5-s time window, during which the infusion of cocaine, the presentation of light cues, and likely the sensation of cocaine’ effects also occurred in addition to the lever-press. It is possible that mice process each of these stimuli through distinct neuronal ensembles that are temporally indistinguishable. In this case, the lever-press ensemble detected in this study is likely a composite of several sub-ensembles, to which D1 and D2 MSNs may differentially contribute.

### Time-dependent changes of the ensemble size

Cocaine self-administration is a learned behavior, involving an initial acquisition of the behavioral sequence, followed by stabilization and maintenance. Over the 11-day procedure, the number of lever-presses increased during the early training days and plateaued over days 7 – 11 (**Fig. 1b**). In parallel, the rotational pattern of the approaching and exiting movements stabilized after days 7-9 (**Fig. 1c-e**). These changes suggest that the cocaine-taking behavioral sequence became more stereotypic and efficient after 7 days of self-administration training, possibly marking a transition from the acquisition to stabilization and maintenance.

During the first few training days, the number of lever-press-responsive neurons increased rapidly (**Fig. 2g**; **3h**). This substantial growth is consistent with the intensive learning and memory encoding associated with acquiring cocaine self-administration. During operant behavioral acquisition, animals face progressively increasing cognitive demands, including adapting to the environment, learning task rules, establishing the stimulus-response association, developing effective behavioral sequence, and incorporating these strategies into memory. To maximize the likelihood of success, excessive recruiting and training are expected (Costa et al., 2004; Brebner et al., 2020; Jáidar et al., 2025), resulting in a large ensemble size.

As training progressed, the mice gradually became familiar with the environment, developed coping strategies, and their operant behavior entered a maintenance phase characterized by stabilized performance. During the transition, NAc neurons continuously received inputs from upstream brain regions related to reward behavior. Repeated co-activation of essential pathways may trigger synaptic remodeling, preferentially strengthening connections representing key behavioral components while weakening irrelevant ones. This re-weighting of synaptic connections may lead to circuit refinement, only keeping a subset of task-specific pathways (Wright and Dong, 2021). As a result, the number of downstream NAc neurons responsive to lever-press gradually decreased, with the turning point on day 7 (**Fig. 2g**; **3h**).

Notably, this broad recruitment followed by refinement during learning is not unique to biological system, as learning in artificial neural networks (ANNs) follows similar principles. During the initial training phase, reducing the size of the network can significantly impair task performance, suggesting that abundant computational resources allocation is essential for effective learning (Tan and Le, 2019). However, after training, certain subnetworks of the original network can perform the task equally well, whereas others contribute little to task execution (Han et al., 2015; Frankle and Carbin, 2019). ANNs are trained by iteratively updating the weights between layers based on inputs, a process analogous to synaptic remodeling during biological learning. This parallel between biological and ANNs suggests that the U-shaped learning curve may represent a general principle shared by both systems.

### Dynamic constitution of the ensemble

The engram theory proposes that memories are coded with discrete ensembles of neurons (Semon, 1921), and these ensembles remain compositionally stable. The hypothesis posits that the same population of neurons forms ensembles during learning, and retrieval reactivates this fixed population (Josselyn et al., 2015; Tonegawa et al., 2018). While this hypothesis is supported by extensive evidence, recent studies show that the composition of such ensembles is dynamic and flexible under certain circumstances (Mau et al., 2020; Zaki and Cai, 2025). Our results revealed that NAc neurons continuously entered and left the lever-press responsive population across 11 days of cocaine self-administration. This flexibility in ensemble composition may represent a key mechanism that allows the brain to update memories by incorporating novel information from ongoing experiences and the ever-changing environment (Mau et al., 2020).

While compositionally flexible, an ensemble can still support stable memory, as explained by competing hypotheses (Zaki and Cai, 2025). One hypothesis suggests a Stable Core, where a small subset of neurons remains consistent, providing a stable foundation for long-term memory (Ziv et al., 2013). Furthermore, the Population Code Stability hypothesis proposes that, while the activity patterns of individual neurons exhibit high variability, the population-level activity pattern remains stable (Driscoll et al., 2017). In this case, neuronal turnover facilitates the incorporation of new task-related information while stability is maintained at the populational level. A third hypothesis proposes that, while the activity of individual neurons is flexible, they collectively form a stable, low-dimensional representation, or manifold, of the key features of the task (Gallego et al., 2017; Perich et al., 2025). Given the limited number of neurons tracked per mouse, our study could not obtain a meaningful representation of a low-dimensional manifold. However, previous studies have revealed that neural manifolds can capture neuronal activity patterns underlying reward behavior (Schall et al., 2024). We therefore hypothesize that such manifold dynamics may represent the underlying mechanism enabling the increasing stability of behavior despite the dynamic composition of the NAc ensemble.

### Concluding remarks

Our study identified a population of neurons within the NAc that was recruited during cocaine self-administration training, forming a potential functional ensemble linked to lever-press behavior. The size of this ensemble exhibited bi-phasic dynamics, reflecting the cognitive dynamics during memory formation versus expression. Ultimately, the stability of drug-taking behavior appears to be encoded not by individual neurons but at the population level, and the development and maintenance of drug-taking behavior might be achieved through an ever-evolving NAc ensemble.

## Acknowledgement

We thank Min Li for technical support. The study was supported by NIH DA040620, DA023206, and DA069868. Cocaine was provided by the NIH NIDA drug supply program.

## Author contributions

LJ, XQ, WJW, BZ, CW, LW, and YD designed the experiments and analyses, and wrote the manuscript. LJ, XQ, JL, and WJW performed experiments and data analysis.

